# Cardiotoxic effects of methamphetamine associated with electrophysiological and epigenetic aberrations in zebrafish

**DOI:** 10.1101/2021.09.16.460189

**Authors:** Jimmy Zhang, Anh H. Nguyen, Lauren Schmiess-Heine, Tai Le, Xing Xia, Michael P.H. Lau, Juhyun Lee, Hung Cao

## Abstract

Long-term methamphetamine (Meth) abuse damages functional and molecular changes in the brain that causes chronic and relapsing disease. In this study, we sought to investigate a relationship between cardiotoxicity and arrhythmia with associated Meth abuse in zebrafish to identify and to understand the adverse cardiac symptoms associated with Meth as well as to assess the applicability of zebrafish as an appropriate model for cardiac-related drug screening studies. Over a two-week duration, zebrafish were first treated with various concentrations of Meth, ranging from 0 to 50 μM. Immediately after treatment, zebrafish underwent electrocardiogram (ECG) measurement for electrophysiological analysis. Results show that a higher incidence of increased heart rate over the duration of the experiment, corroborating with results from previous human case studies involving Meth users. However, abnormalities commonly cited in those same case studies, such as prolongation of QTc, were not significantly presented in obtained ECG recordings. We have also conducted genetic, epigenetic, and histochemical analysis in an attempt to understand the cardiotoxic effects of Meth on zebrafish cardiac function. These results suggested myocardial damage and decrease in gene expression associated with normal physiological function. Finally, this paper provides insights into potential reasons for the apparent discrepancies in our data with prior research as well as an outlook of zebrafish cardiotoxic drug screening studies.

## Introduction

Cardiotoxicity is one of the most adverse consequences of methamphetamine (Meth) abuse, leading to a notable increase of morbidity and mortality (1). Cardiovascular complications are the second leading cause of death in Meth abusers. Cardiotoxicity can appear early in the course of drug use and can cause numerous significant effects, such as pulmonary hypertension, atherosclerosis, cardiac arrhythmias, acute coronary syndrome, and other associated cardiomyopathies (2). Furthermore, a Meth ‘binge’ study to determine long-term effects discovered that Meth decreased the sensitivity of nervous and cardiovascular physiology through successive treatments, implying the potential remodeling of physiological responses through chronic Meth abuse (3). However, data on the mechanism of the appearance of cardiac dysfunction during drug abuse and the susceptibility of long-term development of cardiotoxicity are limited. Therefore, further investigation of Meth abuse and early diagnosis of pre-clinical cardiac dysfunction in zebrafish, a relevant model for human cardiac studies, is essential in order to understand Meth-induced cardiac toxicity.

Meth is a sympathomimetic amine with a range of adverse effects upon multiple organ systems. Based around a phenylethylamine core, Meth and its analog, *d*-amphetamine, have high affinity with transporters associated with catecholamine signaling, significantly increasing the number of neurotransmitters such as dopamine and norepinephrine (1). Unlike Meth, *d*-amphetamine has been prescribed as medication to treat neurological disorders such as attention deficit hyperactivity disorder (ADHD) and narcolepsy (4). A possible reason for limited legal Meth use is the addition of the *N*-methyl group compared to amphetamine, which has been shown to confer better penetration through the blood-brain barrier for Meth, leading to stronger and more addictive responses (5). Meth has been shown to induce heightened catecholamine response by promoting catecholamine release, preventing their reuptake, and destabilizing their levels (6, 7). Thus, Meth is responsible for numerous neurotoxic symptoms (7). Given the elucidation of the direct mechanism of Meth on neurological response, the major focus in researching treatments for Meth-related abuse has been associated with neurological modulation. Therefore, less attention was given to researching the direct mechanism of Meth in other physiological systems, such as the cardiovascular system.

Despite the prevailing issue of Meth abuse, studies have shown that cardiac pathology induced by Meth can be attenuated and even reversed through the discontinuation of Meth use and the initiation of subsequent treatment (8). A study in rats regarding the administration of Meth and eventual withdrawal revealed that the rats were able to recover from myocardial pathologic symptoms such as atrophy, fibrosis, and edema starting from 3 weeks after discontinued Meth administration (9). A human case study indicated that attenuation of Meth use and subsequent therapy led to recovery from ventricular hypertrophy and electrocardiogram (ECG) ST deviations (10). Evidence of recovery from Meth abuse has given promise to future potential treatments, but it also necessitates the research to understand the specific mechanism behind Meth-induced cardiovascular pathologies.

Zebrafish has recently emerged as an efficient low-cost animal research model that has been used for high-throughput phenotype-driven drug screenings for new insights into chemical toxicity due to similar drug metabolism and genetic homology (11). Although current research has not fully proven whether the single ventricular heart of zebrafish is comparable to the more complex ventricular conduction system found in higher vertebrates, zebrafish have been proposed as a versatile model system for biological studies due similarities with humans pertaining to cardiac physiology. Our group has reported the use of *in vivo* surface ECG in adult zebrafish for monitoring irregular heartbeats and QT prolongation (12). Here, we further demonstrated ECG screening in adult zebrafish as a powerful tool for cross-sectional and longitudinal studies for Meth-induced cardiac toxicity. Specifically, we established techniques that allow for the accurate and reproducible detection of irregular arrhythmic symptoms, enabling early ECG-based detection of cardiac dysfunction for drug abusers, further defining ECG as a suitable benchmark test for the extent of Meth-induced cardiotoxicity.

## Materials and methods

### Zebrafish husbandry and experimental preparation

Wild-type zebrafish were housed in a custom-built circulating fish rack system. The water environment was maintained at 28°C, ∼pH 7.0, and was checked at least once daily. The system is equipped with four filters, a UV, carbon, and two particle filters. Zebrafish were kept under the 14:10 hour light/dark cycle. All zebrafish used in this study were wild-type strain obtained from the UC Los Angeles aquatic facility and were approximately 1 year old at the onset of the experiment. Prior to Meth treatment, zebrafish underwent open chest surgery to improve subsequent ECG signal acquisition (12). Under a stereo microscope, scales (above the coelomic cavity and the posterior site above the tail) were removed with forceps, exposing flesh, to allow more direct electrode contact. A small incision, ∼2-3 mm, was made on the ventral surface of the fish above the heart. The incision cut through the chest wall and the heart was visible afterward. The fish were recovered in fresh fish water for a few minutes. The incision was not closed via suture, staple, clips, or glue as chest wall and scales have been observed to regrow within 4 weeks. Fish will regrow scales and recover fully within 4 weeks. The 4 zebrafish groups were then housed in separate tanks throughout the duration of the study. Fish were checked every day until the experiment has concluded, to see whether the animal was slow or showed some abnormal activities (*i*.*e*., erratic swimming, strained breathing, bloating). In those cases, the fish would be euthanized in accordance to UC-Irvine’s Institutional Animal Care and Use Committee (IACUC) guidelines (Protocol # AUP-21-066).

### Drug preparation and Meth treatment

The various concentrations (0, 25, 40, and 50 μM) of Meth solutions for treatment were prepared by mixing the specified amount of Meth stock into water obtained from the fish rack system on the day of treatment. The Meth stock (1 mg/mL) was obtained from Sigma-Aldrich (MDL MFCD00056130). The solution (∼10 mL) was then placed in a small custom PDMS chamber suitable for housing one zebrafish. Each zebrafish was treated in the designated concentration of Meth solution for 20 minutes before ECG recording.

### ECG recording procedure and instrumental setup

ECG was obtained using the instrumental setup depicted in **Fig 1a**. Prior to placing the fish in the designated zebrafish station, the fish was first anesthetized in tricaine (200 mg/L) for approximately 3 minutes until the fish became unresponsive to external stimulus. Then, the fish was placed ventral side up in a precut crevice in the middle of the sponge. The sponge was then placed on a glass platform with the pin electrodes as positioned in **Fig 1a**, where the green working electrode was placed near the open incision on the chest while the yellow reference electrode was placed near the lower abdomen. Each fish underwent recording for approximately 1 minute before placing in regular fish water for recovery from anesthesia. Treatment concentrations and durations were approved by University of California-Irvine’s IACUC.

**Fig. 1.**
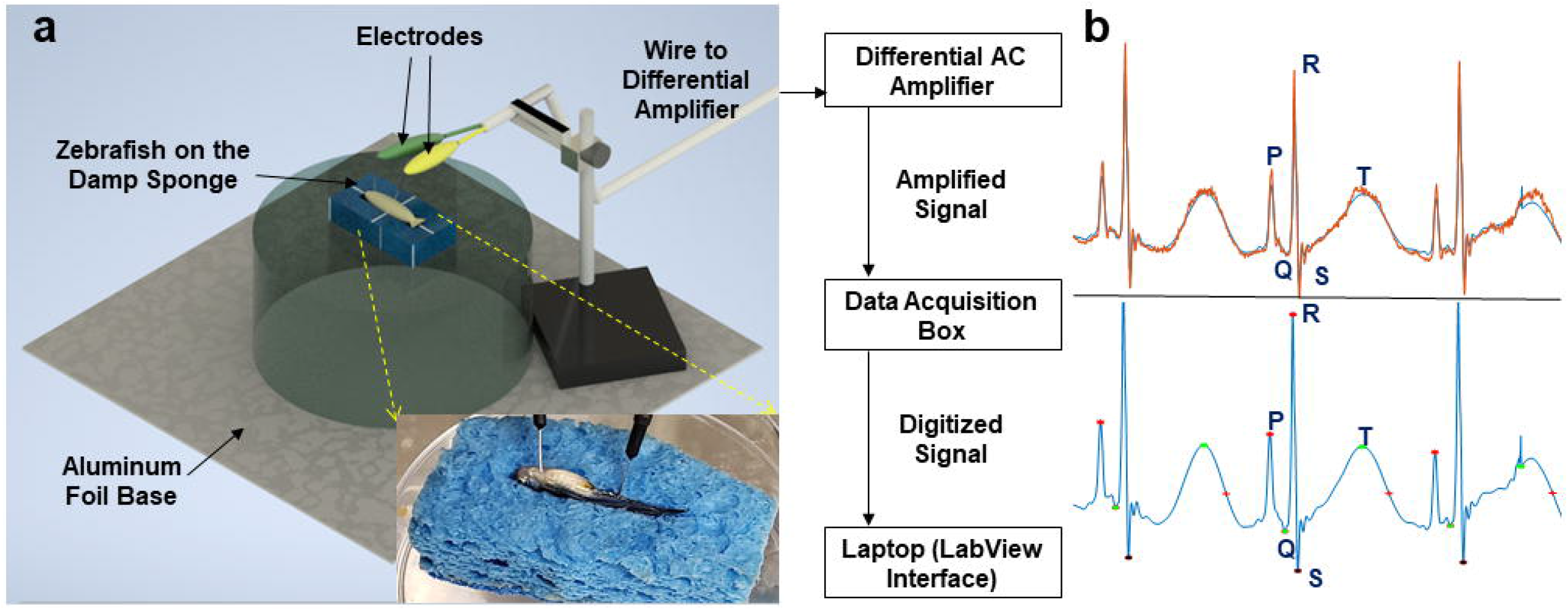
Zebrafish ECG Setup. (a) The figure on the left shows the general layout of zebrafish and electrodes during ECG recording. The working electrode is shown in green and contacts the chest cavity. The reference electrode is shown in yellow and contacts near the tail. This electrode setup connects with instruments outlined in the block diagram on the right, where signals are processed and displayed on the laptop. The islet provides a closer view of the positioning of electrodes on the zebrafish during recording. (b) Representations of ECG signal processing with labeled waveforms utilizing custom Matlab software. The top displays the orange raw signal, while the bottom displays the blue processed signal.

The components of instrumental setup consisted of the following (**Fig 1a**). The pin electrodes were derived from isolating ends of jumper wires (Austor Part AMA-18-580) and stripping the outer insulator layer of the wire about 2 cm from the metal tip of the wires. The exposed copper wire underneath was then soldered to improve signal integrity and electrode longevity. The pin electrodes were attached to alligator clips at the end of a cable wire leading to the differential AC amplifier (A-M Systems Model 1700). The connecting cable was wrapped with aluminum foil and grounded to reduce external electrical noise. The signal from the zebrafish underwent 10000x amplification before undergoing digitalization with the data acquisition box (National Instruments Model USB-6001). The signal was then transmitted to a laptop (Dell Latitude E5470), where the processed ECG signal was displayed using the LabView interface. Examples of ECG recordings are shown in **Fig 1b**.

### ECG data collection and analysis

ECG signals were saved as a text file using the LabView interface. The ECG was subsequently analyzed using a custom-designed MATLAB software to determine the positions of ECG waves for each recording(13). Briefly, the MATLAB m-file first loaded and plotted the ECG data from the output text file from LabView. The m-file then determined locations of peaks corresponding to R waves via local maximum detection. After a denoising step, the P waves, QRS complexes, and T waves were then determined based on predetermined parameters and deviations from the R waves. Finally, relevant statistics such as the QTc duration based on the Bazett formula and heart rate (HR) were calculated and tabulated in an output Excel file corresponding to each ECG data file (14). Prior to subsequent analysis, each data file was reviewed for accurate ECG wave detection by determining that the locations of P, Q, R, S, and T waves depicted by the software conform to the corresponding peaks/troughs on the ECG. Bazett formula: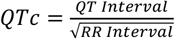

### RT-PCR analysis

Zebrafish were euthanized and homogenized. RNA was successfully isolated following the TRIzol^®^ reagent protocol (Invitrogen). Subsequent first strand synthesis of cDNA was conducted following the protocol of M-MLV reverse transcriptase (Promega). qPCR was conducted with SYBR-Green reagent and analyzed with the Quantstudio Real-Time PCR System. DNA primers used for qPCR are shown in **Table 1**.

**Table 1.**
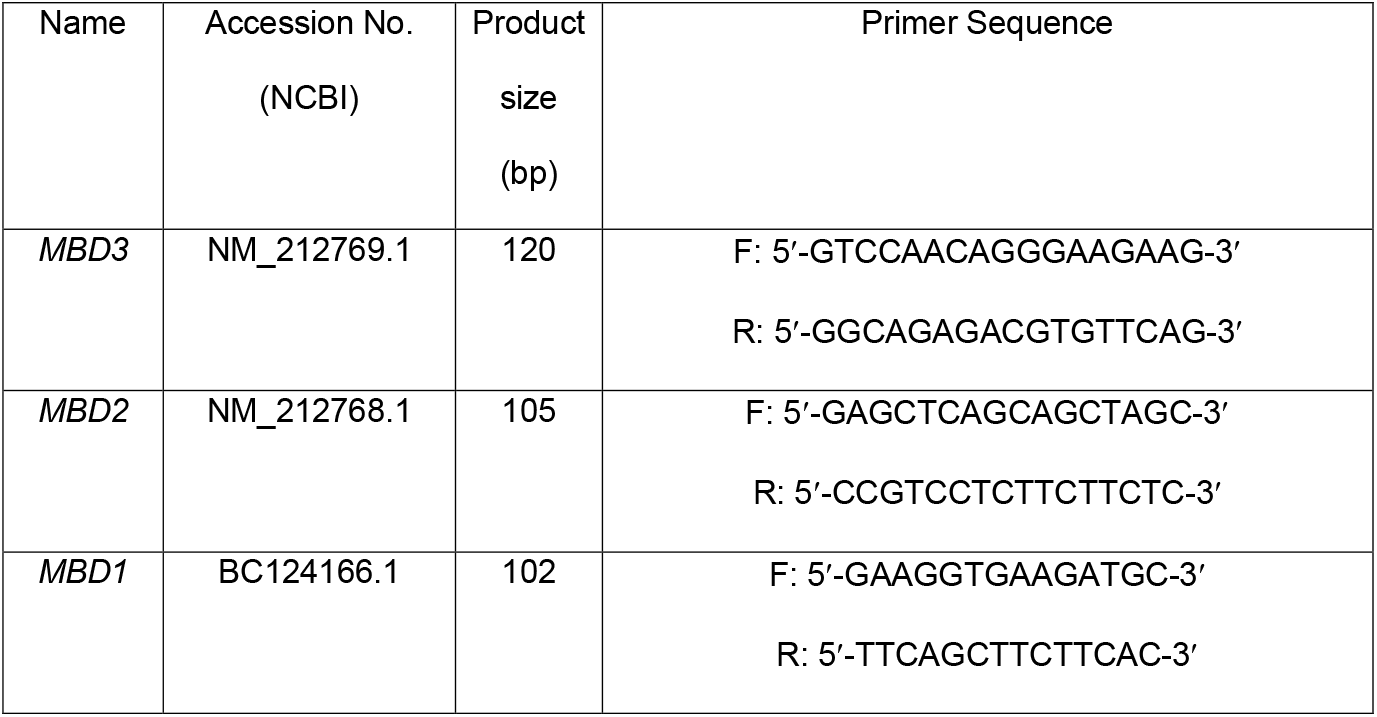
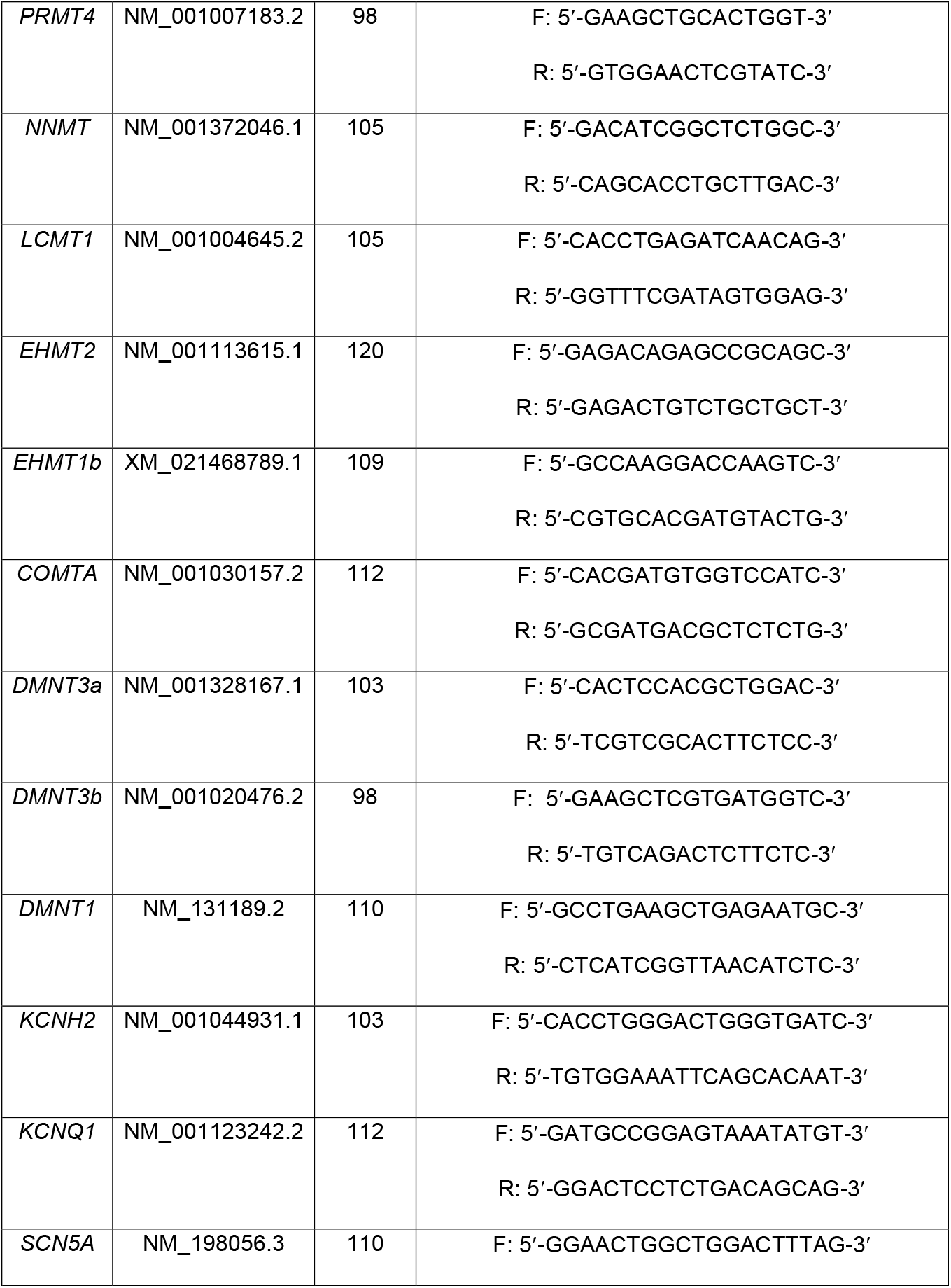
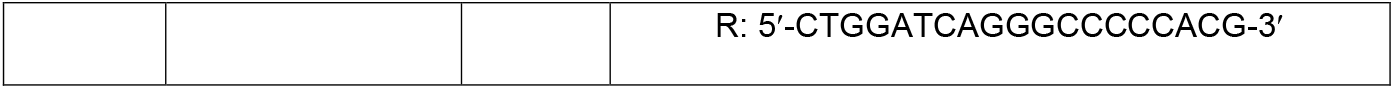
List of primers used for RT-PCR.

### Protein extraction and dot blot analysis

After treatment of zebrafish with Meth concentrations of 25, 40, and 50 μM, the zebrafish were euthanized and homogenized. Their proteins were extracted with the TRIzol^®^ reagent and protocol (Invitrogen). In brief, whole zebrafish (∼400 mg) are gently pipetted in a 1.5 mL Eppendorf (EP) tube. The zebrafish tissues were homogenized by liquid nitrogen before being incubated in 1 mL TRIzol^®^ on ice for 5 min. The extract was centrifuged at 13,000 rpm at 4°C for 45 min and the supernatant was collected into a fresh 1.5 mL EP tube. 200 μL of Chloroform was added and mixed thoroughly, left the tube on ice for 5 min, and centrifuged at 13,000 rpm at 4°C for 15 min. After removing aqueous phase for RNA and precipitating DNA by ethanol, proteins were precipitated by adding 1.5 mL isopropanol, mix thoroughly and incubate for 10 min at room temperature. The extract was centrifuged at 12,000 rpm at 4°C for 10 min, supernatant was discarded. 2 mL of 0.3 M guanidine hydrochloride solution was added and incubate for 20 min at room temperature. The sample was centrifuged at 12,000 rpm at 4°C for 5 min, discard the supernatant. The protein pellet was washed with 2 mL of ethanol and dried for 5 min. The protein pellet was resolubilized with 1mL of 1% SDS solution, the supernatant was collected after centrifuge at 12,000 rpm at 4°C for 10 min, ready to use for subsequent protein analysis. Extracted proteins were dissolved in 1% SDS. The protein solution was then dropped on a nitrocellulose membrane (GE Healthcare Protran^®^ BA85) to form 2 μL and 5 μL dots for each concentration group. After 1 hour of incubation, the membrane was then blocked with TBST including 1% BSA for 1 hour, followed by an overnight incubation with the anti-Meth antibody (mouse, 10M25A Fitzgerald). After washing the membrane with TBST, the membrane was then incubated with HRP conjugated anti-mouse IgG (R&D Systems) for 4 hours. After additional washing, the membrane was then incubated in Pierce™ ECL Western Blotting Substrate (Thermo Scientific) and imaged for chemiluminescence.

### Histochemical staining with Masson’s trichrome stain

After treatment of Meth, zebrafish hearts were isolated as described in previous literature (15). Briefly, the fish were anesthetized by tricaine before undergoing chest incisions. The heart was then located and excised, which were then placed in a solution of perfusion buffer (10 mM HEPES, 30 mM taurine, 5.5 mM glucose and 10 mM BDM in 1x PBS solution). After excision of all hearts, they were subsequently fixed by 4% formaldehyde and cryo-sectioned with a cryostat. The tissue slices were placed onto frosted microscope slides (Thermofisher Scientific) and underwent Masson’s Trichrome staining (following the provided protocol from American Master Tech, TKMTR2). The slides were observed under the microscope for subsequent analysis of collagen and myocardium distribution for cardiac injury analysis.

### Antibodies for immunostaining and ELISA analysis

Protein extraction by TRIzol^®^ reagent was conducted prior to ELISA. Heart tissue samples were fixed and sectioned with the aforementioned procedure prior to immunostaining. The antibodies used for immunostaining and ELISA were mouse anti-Meth antibody (mouse, 10M25A Fitzgerald), anti-MBD2 (ab45027; Abcam), anti-DNMT1 antibody (mouse, ab13537; Abcam) and the secondary antibodies used were goat anti-mouse IgG-HRP (ab6789; Abcam), goat anti-mouse IgG H&L (Alexa Fluor® 488) (ab150113, Abcam).

### Statistical analysis

All parameters derived from data analysis (*i*.*e*., QTc, HR) were averaged within each experimental group. Statistical significance was determined by the one-way ANOVA test between experimental groups with significance level *p<*0.05.

## Results

### Significant dose-response increase in heart rate but not in QTc from Meth treatment

Zebrafish has been a favorable model for drug screening experiments, including Meth. Indeed, most studies regarding zebrafish Meth treatment concern the neuropsychological effects of Meth, owing to the ability of Meth to enhance catecholamine neurotransmitters (16, 17). Studies of zebrafish cardiotoxicity in response to Meth administration are scarcer. Fang *et al*. conducted a Meth cardiotoxicity study by treating 48 hpf (hours post fertilization) zebrafish embryos with up to 24 hours of Meth administration, documenting decreased heart rate and abnormal morphological changes such as edema and hemorrhage (18). To this end, we sought to verify and expand on their results through our methodology in ECG acquisition, as arrhythmia was a prominent cardiotoxic symptom in Meth abusers (1). Our lab previously developed modalities in acquiring ECG from adult zebrafish and successfully identified abnormal ECG characteristics (12). Using an ECG setup designed in our lab (**Fig 1**), we acquired ECG signals over a two-week period. Each experimental group is treated with the designated Meth concentration for 20 minutes for 3 instances per week, and their ECG was subsequently acquired after treatment. The signals were then processed to eliminate external noise, and parameters such as heart rate and QTc were then quantified. **Fig 2a** comprises representative ECG figures obtained from our ECG system, including ECG diagrams obtained before Meth administration as well as 7 days and 14 days after first administration. **Fig 2b** is a representation of a pathogenic progression of ECG signals due to Meth treatment. The evident increase of heart rate and sporadic signs of QTc prolongation were seen after the first and second weeks of treatment. Through our analysis of ECG recordings of zebrafish treated at varying concentrations of Meth over the period of 2 weeks, a significant increase in heart rate was noted for all zebrafish groups treated with Meth compared to zebrafish not treated with Meth (**Fig 2a**), which is consistent with previous studies of Meth users displaying monomorphic ventricular tachycardia (19). However, our data indicated that significant QTc prolongation did not occur due to Meth treatment (**Fig 3**). This seemingly contradicts with results obtained from previous human medical case studies, as those cases reported significant signs of QTc prolongation in Meth users (20, 21). Comparison between heart rate and QTc for individual measurements (**Fig 3a**) revealed that there was no significant correlation between heart rate and QTc. Furthermore, heart rate tended to display a wider range of values than for QTc across treatment groups, supporting the previous analysis that Meth induces a more significant change in heart rate than in QTc. Heart rate values for each treatment group were more delineated, as values from the control group were colocalized at the lower range of heart rates (mean = 87.4 bpm), whereas values from the 50 μM treatment group were colocalized at the higher range (mean = 94.6 bpm). QTc results did not display the same discrimination as in heart rate, as values were congregated regardless of experimental group. This was corroborated by measuring the variation in heart rate in each group, as the variations remain relatively similar regardless of experimental treatment. While some outliers populated the high extreme values of QTc, the majority of QTc values were congregated.

**Fig 2.**
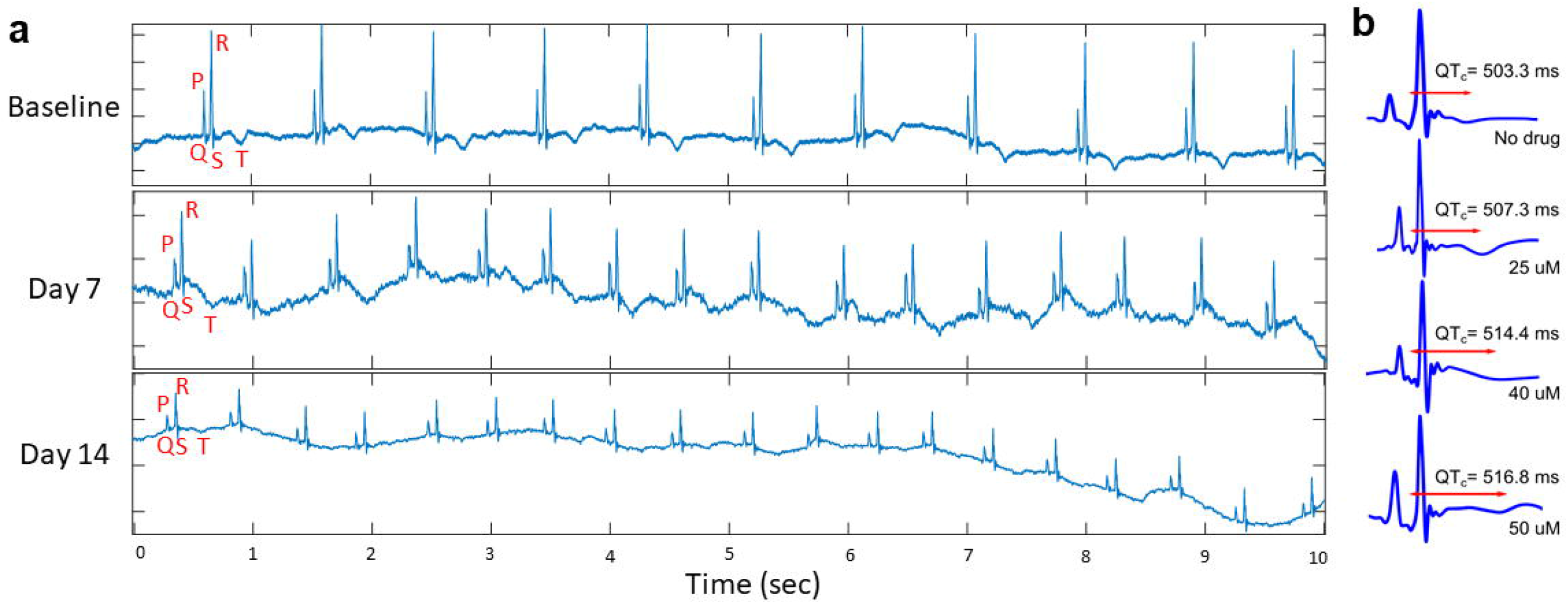
Analysis of ECG before and after treatment. (**a**) This representative figure indicates the ECG signals before treatment, labeled as ‘Baseline’, as well as signals obtained at 7 and 14 days after the onset of methamphetamine treatment. ECG waveforms were labeled for the initial ECG wave. The progression of ECG signals depicts an increase in heart rate after onset of treatment, with sporadic cases of QTc prolongation. (**b**) Representation of average QTc measured for each experimental group across the duration of treatment, indicating an increase of QTc, but at an insignificant level.

**Fig 3.**
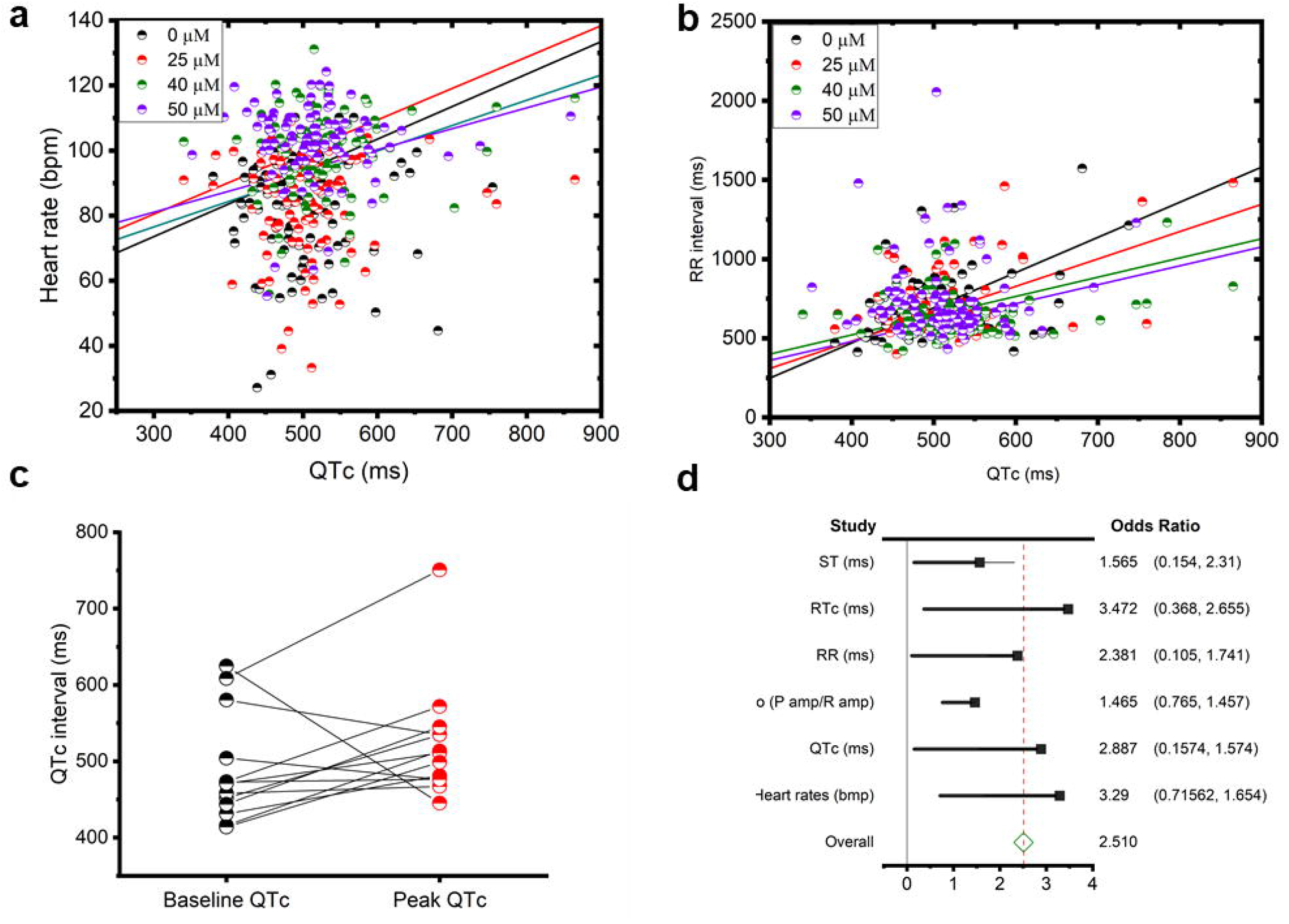
Comparison of QTc and HR across experimental Groups for significance. Correlations of heart rate and QTc (**a**) and correlation of RR interval and QTc (**b**), indicating a general positive correlation between heart rate and QTc. (**c**) Correlation of baseline QTc values obtained before methamphetamine treatment with peak QTc measurements obtained during the course of the treatment, suggesting a general prolongation of QTc. (**d**) Odds analysis of ECG parameters, indicating the likelihood of significant changes before and after methamphetamine treatment. As shown, RTc, QTc, and heart rate indicate significant changes with OR>2.5.

### Lack of significant differences in ECG characteristics between treatment groups

While significant differences can be found for heart rate between zebrafish treated and not treated with Meth, other characteristics, including ST intervals and RR intervals, do not show significant correlation. According to the odds analysis shown in **Fig 3d**, only QTc, heart rate, and RTc have shown significant differences between Meth-treated and untreated groups (OR > 2.5). A closer inspection of the odds ratios indicate that the significance was fairly weak for RTc and QTc, which was corroborated by T-tests conducted above. Other parameters that were studied, such as ST interval, have an odds ratio below the significant value.

### Sporadic dysregulation of ST interval and other ECG characteristics after Meth treatment

While certain ECG parameters do not display significance between treatment groups, an analysis into the variability of these parameters revealed interesting results. Notably, the variability in ST interval increases at higher concentrations (**Fig 4**). While the mean ST interval calculated was not significantly different at all Meth concentrations, the sporadic presence of increasing ST interval duration as Meth concentration increases indicated the propensity of ST interval to vary after Meth dosage. This pattern was not seen in QTc and heart rate, as the variation of both parameters remain similar throughout all treatment groups. Interestingly, RTc also displayed a similar variation to ST, but the difference in RTc may be attributed to ST. While the significance of a prolonged ST interval is not well understood, a previous manuscript documented that ST interval prolongation could be associated with left ventricular hypertrophy (22). This could be explained by realizing that the increase in ventricular volume potentially increases the distance that the Purkinje fibers have to cover for both ventricular depolarization and repolarization, increasing the ST interval.

**Fig. 4.**
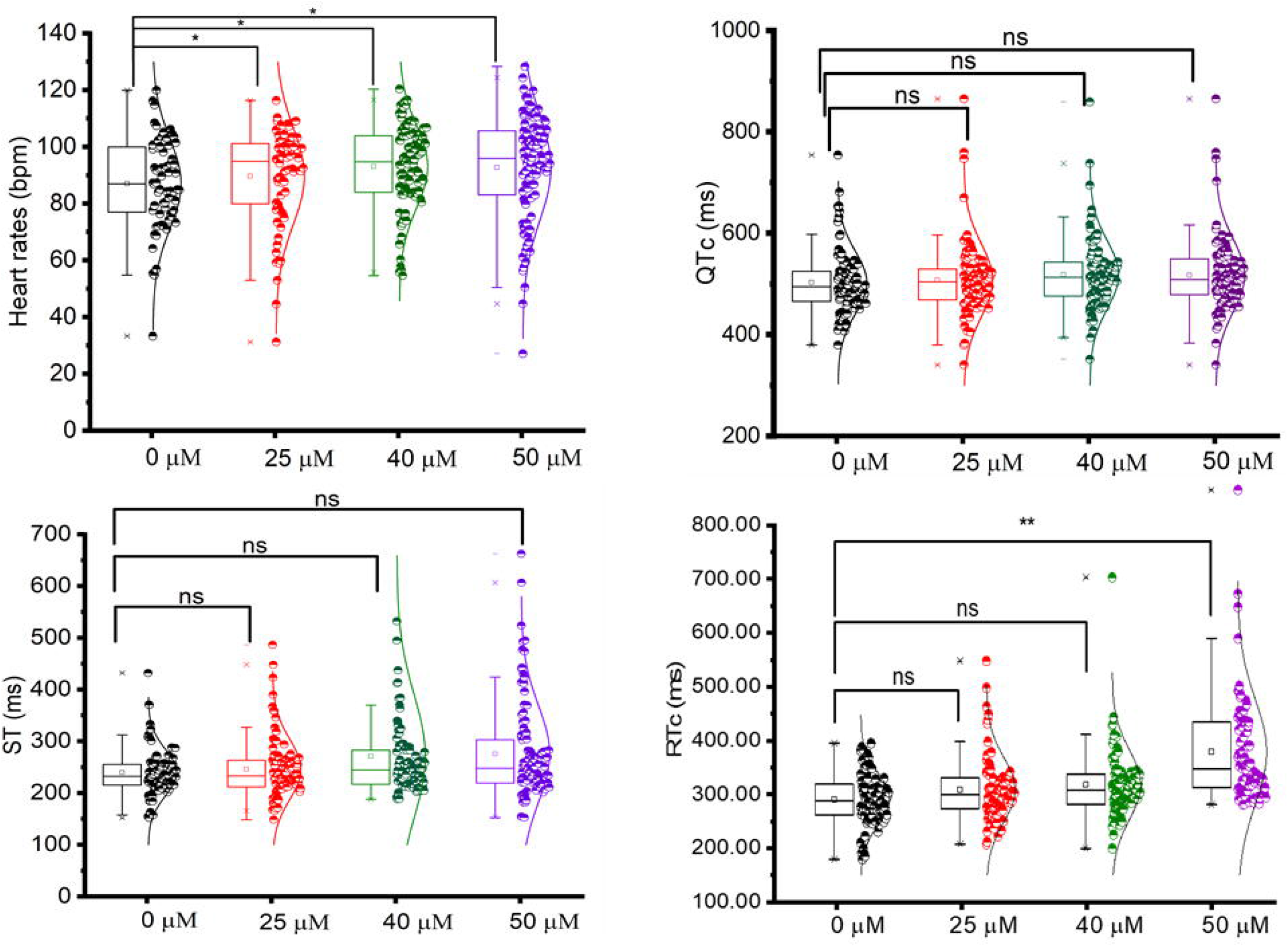
Analysis of specific ECG parameters between experimental groups. Aggregate measurements were plotted for each experimental group for comparative analysis. Based on significance testing, heart rate exhibited significant increases across all experimental groups, supporting that methamphetamine induces increased heart rate in zebrafish. Other parameters (QTc, RTc, and ST) displayed little to no significance across experimental groups. * indicates *p* < 0.05. *ns* indicates no significance.

### Ion channel dysfunction associated with Meth treatment

While we did not observe significant changes in ECG signals between Meth treated and untreated groups aside from heart rate, we wanted to determine if Meth has induced cardiotoxic damage in cardiac tissue. Through our dot blot analysis to detect Meth within zebrafish tissue, we were able to determine that Meth does retain within zebrafish after treatment in a dose-response manner (**Fig 5f**). We chose to investigate and genetic and histological consequences of Meth treatment in zebrafish. To analyze molecular roles of Meth in inducing cardiotoxicity, we conducted RT-PCR of several genes associated with ion channels. These potassium and sodium ion channel genes (*KCNH2, KCNQ1, SCN5A*) have been selected due to their arrhythmogenic property seen in arrhythmic disorders (23). While the potassium ion channel genes *KCNH2* and *KCNQ1* indicated no significant expression changes, *SCN5A* displayed significant upregulation in the Meth treated groups compared to the untreated group (**Fig. 5a**). *SCN5A* corresponds to the sodium channel protein Na_v_1.5. Through previous studies involving the characterization of gain-of-function *SCN5A* mutations, upregulation of the sodium ion channel is known to be a factor in several arrhythmic diseases, including Brugada and long QT syndromes, as well as cardiovascular diseases such as cardiac hypertrophy and heart failure (24, 25). Therefore, Meth treatment had been demonstrated to potentially induce arrhythmogenic consequences directly due to dysregulation of certain ion channel expressions and functions.

**Fig. 5.**
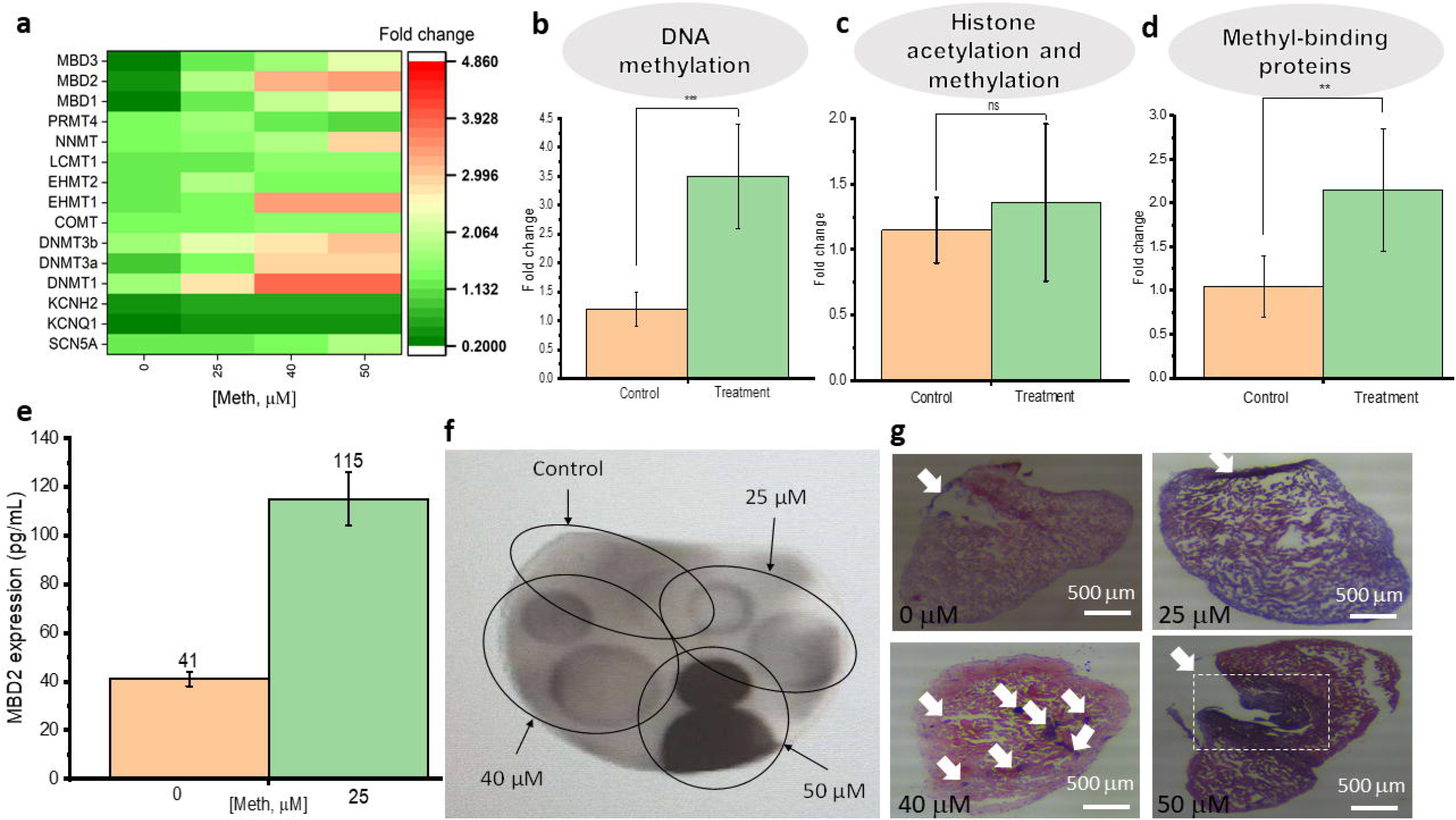
Genetic Analysis of Methamphetamine Treatment on Zebrafish. (a) The degree of expression of genetic and epigenetic genes in comparison between experimental groups. Results indicate that MBD1-3, NNMT, EHMT1, DNMT1, and DNMT3A-B exhibit the greatest progressive fold change in expression as methamphetamine dosage treatment increases. SCN5A, a sodium channel gene common in arrhythmic syndromes, has also been upregulated by methamphetamine treatment. (b-d) Quantification of expression changes based on functionality of epigenetic proteins (categorized as DNA methylation, histone methylation and acetylation, and methyl-binding proteins). The protein expression across all categories is increased in methamphetamine treated group (25 μM), with the difference being significant in DNA methylation proteins and methyl-binding proteins. In the panel of genes tested, 29% of genes are associated with DNA methylation, 58% with histone acetylation and methylation, and 13% with methyl-binding proteins. (e) Comparison of protein expression of MBD2, a methyl-binding protein, producing the most pronounced difference in expression between control and 25 μM treated group. (f) Dot blot diagram of methamphetamine detection retained in zebrafish tissue, indicating that methamphetamine does retain in zebrafish tissue in a dose-dependent response. (g) Representative trichrome-stained cardiac sections from zebrafish treated at various concentrations of methamphetamine. Red and pink stains from Biebrich scarlet-acid fuchsin dye indicate cardiac muscle tissue. Blue stains from the aniline blue dye indicate collagen fibers from scar tissue. White arrows and dashed box indicate the locations of collagen deposition in the cardiac tissue. Overall, cardiac tissue derived from zebrafish treated at higher concentrations (40 μM and 50 μM) displayed more tissue injury due to higher collagen deposition than lower concentrations. Scale bar = 500 μm. * indicates significance with p < 0.05. ns indicates no significance.

### Transcriptional repression as an epigenetic change due to Meth treatment

To further analyze the genetic consequences of Meth treatment, we determined the expression changes of epigenetic factors in zebrafish after Meth treatment. From our analysis, the most noteworthy result shown was the expression change in *MBD2*. Zebrafish treated with 25 μM Meth resulted in an approximately 3-fold change in *MBD2* compared to those in the control group (**Fig 5e**). *MBD2* corresponds a transcriptional repressor that specifically binds to CpG sites formed from DNA methylation, so the upregulation of *MBD2* is an implication of long-term repression of genes. Analysis of other potential epigenetic factors potentially affected by Meth treatment yielded similar results. In the diagram depicted in **Fig 5a**, the factors that have underwent a significant fold change (by at least 2 fold) include *MBD1-3, NNMT, EHMT1, DNMT1*, and *DNMT3A-B*. These factors are mainly involved in the various forms of methylation, including DNA methylation and histone methylation. These results were also supported through our quantification of the expression of DNA methylation, histone modifications, and methyl-binding proteins between the control group and the 25 μM treatment group (**Fig 5b-d**). As demonstrated in a previous study analyzing epigenetic influences in *SCN5A* expression, *SCN5A* dysregulation was associated with epigenetic regulations (26). Thus, the upregulation of DNA methylation could potentially be linked with the dysregulation of pertinent ion channel genes such as *SCN5A*. Interestingly, our results yielded a significant difference between the control and treatment groups for DNA methylation and methyl-binding proteins, but not for histone modifications. Histone modifications are generally associated with short-term, reversible forms of genetic repression, whereas DNA methylation is correlated with long-term and stable genetic repression (27). Masson’s trichrome stains were performed in cardiac cryosections in which myocardial fibrosis are indicated in blue at 40 and 50 μM of Meth, compared to that of control (0 μM) and 25 μM of Meth (**Fig. 5h**). These results were also corroborated through our histological analysis, by specific antibodies in fluorescent immunohistochemistry (**Fig 6**), further supporting that Meth induces an upregulation in both DNA methylation and subsequent transcriptional repression.

**Fig. 6.**
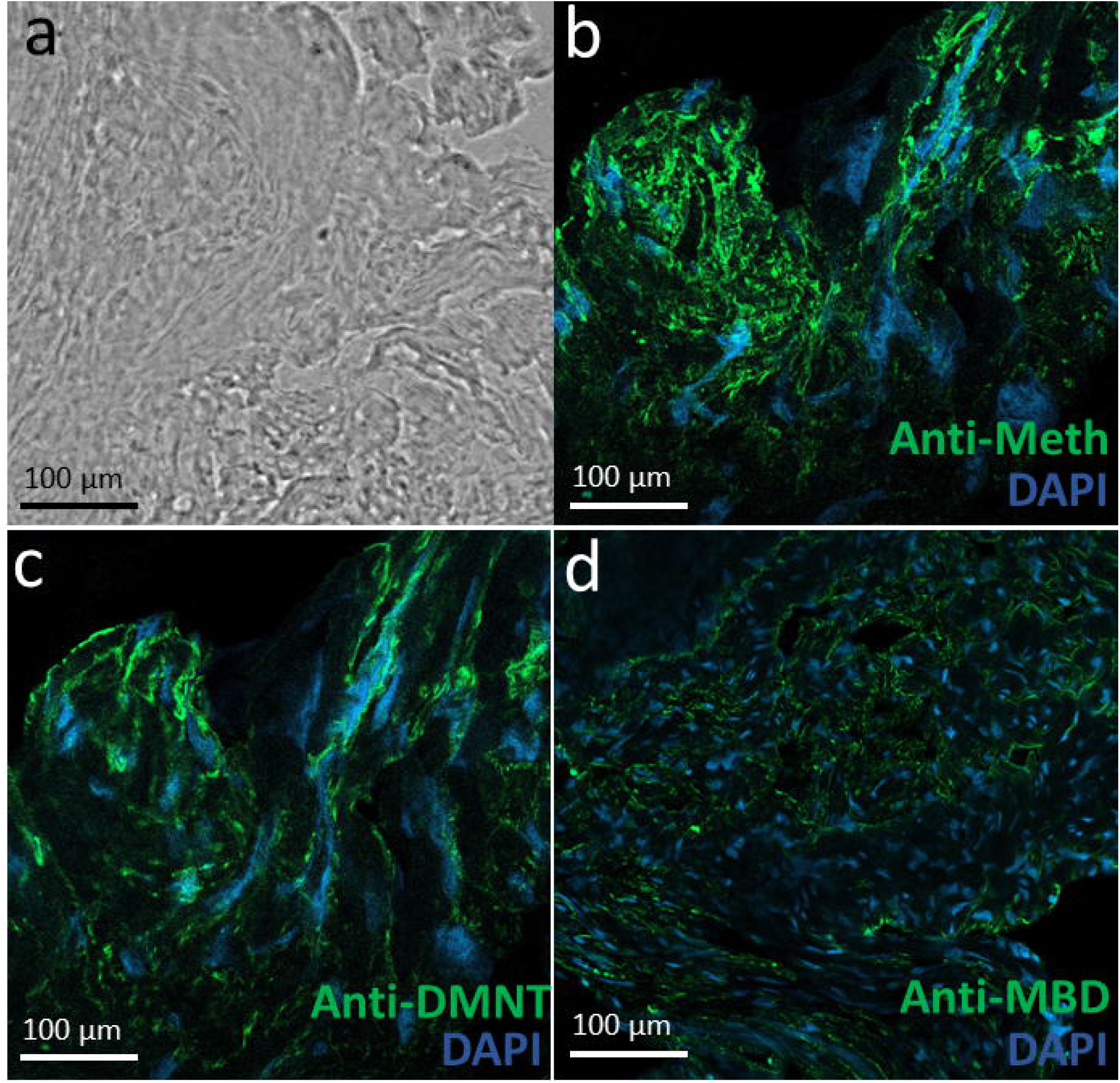
Representative Immunostaining of Methamphetamine and Epigenetic Proteins. (a) Brightfield image of zebrafish heart tissue section. (b-d) Immunostaining of methamphetamine (b), DMNT (c), and MBD (d) within cardiac tissue derived from methamphetamine-treated zebrafish. Antibodies were conjugated with Alexa Fluor 488 to indicate presence of proteins with green fluorescence. The immunostaining figures indicate the retention of methamphetamine after treatment, as well as the upregulation of DMNT, a DNA methylation protein, and MBD, a transcriptional repressor in response to DNA methylation. Scale bar = 100 μm.

## Discussion

The Meth epidemic continues to fester worldwide, and cardiovascular diseases are the leading cause of death for Meth abusers. Utilizing animal models to study cardiovascular associated mechanisms could be critical in devising treatments for Meth associated diseases. The zebrafish is an excellent model for drug screening studies due to high fecundity, low maintenance, and similar genetic homology to that in humans. As the zebrafish model is constantly evolving, studies have continued to delineate the applicability of the zebrafish in human medical research. During the initial conception of this study, we have sought to 1. Establish the zebrafish model as an adequate model of drug screening for cardiotoxic effects, and 2. Elucidate the role of Meth in inducing long-term cardiotoxic effects. Utilizing our custom-built zebrafish ECG acquisition system in our lab, we were able to acquire ECG from zebrafish during the two-week treatment period with Meth. Based on our results, we determined that heart rate has increased significantly during the two-week period due to Meth treatment through comparison of heart rate between zebrafish treated with Meth and zebrafish that were treated in normal fish water. Additionally, we conducted an analysis on possible gene targets of Meth that are associated with cardiotoxicity and discovered that *SCN5A*, a cardiac sodium ion channel gene commonly associated with arrhythmogenic syndromes, was significantly upregulated due to Meth treatment. Previous studies have shown that deregulation of these genes have resulted in deterioration in cardiac physiology, including the presence of prolonged QTc (28). These results were consistent with previous human case studies (1). Additionally, we have conducted some experiments to analyze possible Meth-induced epigenetic changes, and results indicated that Meth intake was correlated with an increased amount of DNA methylation and associated gene repression. Similar results have been documented for Meth-induced addiction studies in other animal models (29, 30). While hypermethylation could possibly be associated with the upregulation of ion channel genes and the subsequent dysregulation of cardiac rhythm, the exact mechanism linking these epigenetic and physiological changes remains unknown. In the future, experiments should be conducted to determine the gene targets of hypermethylation. Nevertheless, our results have indicated the propensity of Meth to induce cardiotoxicity in zebrafish through genetic and epigenetic means.

While some associations have been determined between Meth treatment and pathologic symptoms shown on ECG in zebrafish, the significance is not very evident. For example, there is currently no explanation on why a dose-response increase in Meth was not seen in certain relevant parameters in ECG. While the dose-response was evident when measuring heart rate, the determination of significance in other parameters was less apparent, such as the presence of QTc changes in response to variations of treated Meth concentrations. QTc prolongation has long been regarded as a prominent symptom in Meth users, and in some isolated cases in this study, QTc prolongation can be seen on ECG signals. However, statistical analysis indicated that there were no significant QTc changes due to Meth treatment. Despite the lack of significance, Meth was confirmed to successfully retain within the zebrafish after treatment, as verified by our dot blot experiment. Meth retention in zebrafish was also demonstrated in a previous zebrafish study (31). We surmise that Meth might pose an antagonizing interaction with tricaine, the anesthetic agent used to acquire ECG. Both Meth and tricaine have opposing mechanisms, as Meth is a stimulant while tricaine is an established anesthetic, known for preventing action potential firing by blocking voltage-gated sodium channels (32). Tricaine has also been shown to prolong the Q-T interval as well, which could also confound zebrafish ECG results (33). Most zebrafish Meth studies focused on the neuropsychological effects of Meth, which were conducted using behavioral tests and without the need for tricaine (34). Therefore, future improvements should be implemented to reduce the effect of tricaine for zebrafish cardiotoxic studies, as tricaine could introduce confounding circumstances, especially when testing psychostimulants on zebrafish. Future studies should also seek to explain the effect of these chemical entities on ion channel function through analysis of sodium and calcium transients for zebrafish. These future experiments would determine the mechanism of Meth in inducing cardiotoxicity as well as bolster the use of zebrafish as a suitable model for cardiotoxic studies.

Additionally, future directions would also include linking neurotransmitter response to cardiovascular abnormalities in zebrafish as well as an investigation on ion channel function in the heart after Meth administration. The zebrafish model has already been utilized in numerous Meth studies, mostly related to behavioral studies due to Meth’s known ability to disrupt dopamine release and reuptake, thus increasing dopamine expression (35). Therefore, it would be intriguing to understand the role of dopamine in Meth-induced cardiotoxicity, as it would explain whether Meth-induced cardiotoxicity is caused by dopamine or through a direct effect from Meth. One consequence of dopamine response is the change in ion channel expression. Studies have shown the modulation of L-type calcium channels by Meth, but it is not fully understood whether Meth alters calcium channel function directly or via dopamine (30, 36, 37). Overall, more future studies should be planned to elucidate the mechanism of Meth-induced cardiotoxicity.

## Acknowledgements

The authors would like to acknowledge the financial support from the NSF CAREER Award #1917105 (H.C.), the NSF #1936519 (J.L. and H.C), and the NIH SBIR grant #R44OD024874 (M.P.H.L and H.C.).

## Notes

### Competing Interest Statement

The authors have declared no competing interest.

